# Northward range expansion of the European green crab *Carcinus maenas* in the SW Atlantic: a synthesis after ~20 years of invasion history

**DOI:** 10.1101/2020.11.04.368761

**Authors:** Mariano E. Malvé, Marcelo M. Rivadeneira, Sandra Gordillo

## Abstract

This study aims at synthesizing the recent invasion history of *Carcinus maenas* in the SW Atlantic (~20 years), particularly the northward expansion, based on available published papers, technical reports, and new field surveys. Our analyses extend the known distribution range northwards ca. 330 km. totaling ~1000 km along the Argentinean coast since its last detection in Nuevo Gulf in 2015. The expansion rate appeared to slow down during the last 15 years (from 115km/yr. to 30 km/yr.) as the species continues moving northwards into the transition zone between the Magellan and Argentinean biogeographic provinces (41°–43°S). In addition, a species distribution model (SDM) is provided at a much finer spatial resolution than previous studies, which accurately foresees suitable areas of invasion in the southern San Jorge Gulf, and predicts a hotspot of invasibility around 40°–33°S° if the invasion continues northward. Potential impacts of *C. maenas* on native species, particularly economically important oyster beds are discussed.

## Introduction

The SW Atlantic (SWA) coast has been less exposed to invasions due to the lower intensity of maritime transportation; however, most coastal ecosystems have already been modified, or are expected to be so in the short term (Orensanz et al. 2002). The European green crab (*Carcinus maenas*) is native to the European Atlantic and northwest Africa, with distribution from Mauritania to Norway (Jamieson et al. 2002), and it is one of the 100 world’s worst invasive species (Lowe et al. 2000).

This species has been successfully introduced into several temperate regions outside of its native geographic range, such as Atlantic and Pacific North America, Australia, South Africa, Japan, and Argentinean Patagonia (Carlton and Cohen 2003), although its ecological impacts appear to vary between populations and regions. The first report of *C. maenas* on the SWA coast was registered in 2001 by Vinuesa (2005) in Comodoro Rivadavia (45.52°S) and Rada Tilly (45.93°S), while Hidalgo et al. (2005) recorded the species in Camarones Bay (44.79°S) in 2003 (Fig. 1b). Barón (2007) also detected the species in 2006–2007 from Camarones Bay (44°S) up to the Puerto Deseado estuary (47.8°S), and found maximum abundances at Mazaredo, in the southern San Jorge Gulf. In 2015, the species was reported at Puerto Madryn (42.46°S) in Nuevo Gulf, thus extending the distribution range 250 km northwards (Torres and González-Pisani 2016). The introduction of *C. maenas* in Argentinean Patagonia might have taken place at the Comodoro Rivadavia harbor (southern Chubut province, Fig. 1b), most probably via ballast waters from an oil vessel, and in successive years the population spread to Camarones Bay by northward larval transport (Hidalgo et al. 2005).

**Fig. 1.**
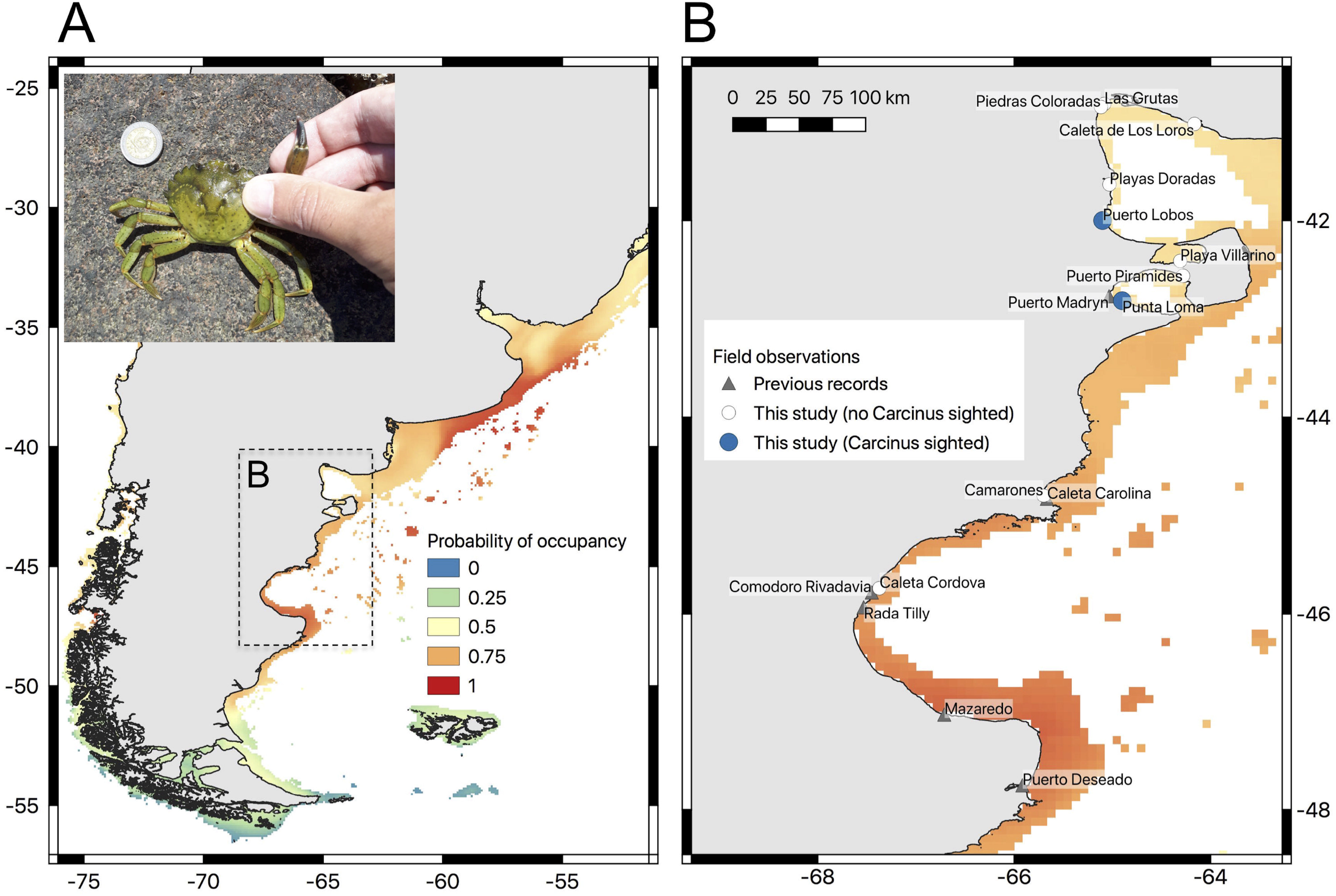
*Carcinus maenas* along the SW Atlantic coast. A) Species distribution model showing the invasion potential of *C. maenas*. The inset shows the specimen collected live in Puerto Lobos, Chubut province, San Matías Gulf, Argentina. B) Sightings of *C. maenas* reported by previous studies and this analysis.

Temperature has been inferred as the most important factor limiting the invasion potential of *C. maenas* (Carlton and Cohen 2003), since *C. maenas* larvae can only develop into adults within a specific temperature range from 10 to 22.5°C (de Rivera et al. 2007). This physiological limitation may explain why this species has colonized far more to the north of its entry point in Argentinean Patagonia than to the south. In fact, the *C. maenas* populations found in Patagonia are the world’s most southern population. Thermogeography models have also identified potential geographic range extensions along many currently invaded coastlines and the potential invasion of countries such as Chile and New Zealand (Compton et al. 2010). However, based on literature analysis and examination of museum specimens, Carlton and Cohen (2003) found that *C. maenas* is absent from climatically suitable regions including the SE Pacific and part of the Atlantic coasts of South America, such as Tierra del Fuego, Chile, the northern border of Peru, and the southern border of Brazil, among other areas around the world.

This study aims at synthesizing the recent invasion history of *Carcinus maenas* in the SW Atlantic (~20 years), based on available published papers, technical reports, and a new finding in the San Matías Gulf (Puerto Lobos, 41.99°S) during December 2019 in a survey of 10 rocky intertidal sites from 41°S–46°S. Range expansion rates were calculated for the first time since its initial detection in the region, and a comparison between rates from other invaded areas around the world is presented. In addition, a species distribution model (SDM) is provided at a much finer spatial resolution than previous studies.

## Materials and methods

### Rate of range expansion

All existing records of *C. maenas* for the SWA (including published papers and technical reports) were used to estimate the geographic expansion range. The northernmost and southernmost record for each year were considered, and estimations were made of the geographic distance from the likely entry point of the invasion at Comodoro Rivadavia (45.46°S). The published year of the first detection for each site was used in the analysis, although the species was probably there some time before it was first reported. This information was complemented with a new field survey carried out in 10 rocky intertidal sites from 41°S–46°S during December 2019. Sampling was carried out during diurnal low tides in the intertidal zone by turning over stones and looking inside tide-pools.

Least-cost distance between sites was estimated taking into account the shape of the coastline and the bathymetric range of the species (0–60 m), thereby providing a more realistic estimation of the distance between two sites. Analyses were carried out using a 1-arc min resolution bathymetry from ETOPO 1 implemented in the library *marmap* (Pante and Simon-Bouhet 2013) in R. A segmented regression was used to estimate temporal changes in the regression slope between distance and year (i.e., expansion rate). Analyses were carried out using the libraries *geosphere* (Hijmans 2007) and *segmented* (Muggeo 2008) in R.

### Species distribution modelling

A species distribution model (SDM) was built based on 27,500 global georeferenced occurrences of *C. maenas* obtained from the OBIS database (www.iobis.org, downloaded on March 17, 2020). Six occurrences reported for the Argentinean coast from previous studies were also included. These occurrences were ‘thinned’, leaving only one single occurrence per 5-arc minute grid cell, in order to ameliorate possible spatial autocorrelation. A final total of 2,091 occurrences were used in further analyses. Occurrences in native and invasive areas were not differentiated (e.g., Compton et al. 2010). Six oceanographic variables (mean SST, SST range, primary productivity, mean salinity, salinity range, pH, and oxygen concentration) were used to model SDM, and were obtained from the BioOracle Database v.2.0 (Assis et al. 2018) at 5 arc-min resolution (ca. 9.2 km). To carry out the SDM, a multi-model ensemble approach was used based on six different algorithms (generalized linear model, generalized additive model, generalized boosted regression, multivariate adaptive regression spline, and random forest) implemented in the library *sdm* in R (Naimi and Araújo 2016). Models were trained using 67% of occurrences, and tested with the remaining 33% across 100 replicates. The ensemble prediction across all models was used to create a map of the potential distribution of *C. maenas* along the SWA, masking the distribution to include only areas < 60 m, which is the maximum known bathymetric distribution of the species.

## Results and Discussion

Our new field survey extends the northward range of expansion of *C. maenas* ca. 330 km, crossing the Valdés Peninsula and entering the San Matías Gulf. One male individual of *C. maenas* was found alive inside a tide pool in the upper intertidal of Puerto Lobos (Chubut province, southern San Matías Gulf) (Fig. 1ab), along with abundant specimens of other native crabs (e.g., *Cyrtograpsus altimanus, Ovalipes trimaculatus*). The single individual of *C. maenas* was photographed alive on site. The carapace measured 53.18 mm in width and 38.85 mm in length (Fig. 1a), which is within the values of carapace width for males previously registered at Puerto Madryn (37.8–89.5 mm, mean = 63.65 ± 36.57 mm, n = 10) by Torres and Pisani-González (2016), and those reported by Hidalgo et al. (2005) at Camarones Bay (51.4–81.4 mm, n = 35). Extensive searches were carried out in an attempt to find more specimens at the intertidal, but these were unsuccessful. Neither were dead individuals found at the supratidal. Although only one specimen was found, the importance of this finding is based on the dispersal of this invasive species into the transition zone between the Magellan and Argentinean biogeographic provinces through the Valdés Peninsula, which is an area declared a world heritage site by UNESCO in 1999.

The current invaded range of *C. maenas* is estimated to be around 1000 km (720 km northwards and 280 km southwards of its likely entry point at Comodoro Rivadavia). The current northward expansion rate of *C. maenas* is 30 km/yr., which is much lower than during the initial phase (115 km/yr.) (Fig. 2). The current expansion rate of *C. maenas* is similar to that estimated for western North America (20–31km/yr., Grosholz and Ruiz, 1996), but is lower than estimates for eastern North America (> 60 km/yr., Grosholz 1996; Jamieson et al. 2002), and higher than estimates for South Africa (16 km/yr., Grosholz and Ruiz 1996) and Australia (1.7 km/yr., Thresher et al. 2003). Fast mean expansion rates (200 km/yr.) have also been documented in San Francisco Bay (western North America) between 1993 and 1999 (Yamada et al. 2000). Rates of range expansion also show great variation across invaded regions, years and sites, and intraregional expansion in some areas has been episodic, with rapid jumps between periods of relative stasis (Grosholz 1996; Thresher et al. 2003). In this context, *C. maenas* appeared to slow down its rate of expansion as the species continues moving northwards into the transition zone between the Magellan and Argentinean biogeographic provinces (41°–43°S, Wieters et al. 2012). Thus, a change in the expansion rate may be expected if this species invades the Argentinean biogeographic province, since the SDM predicts an invasibility hotspot around 40–33°S. This area corresponds to the Buenos Aires coast and was the entry point of the acorn barnacle *Balanus glandula* to the SWA, which has invaded most of the rocky shores of Argentina at a high rate of expansion (Schwindt 2007).

**Fig. 2.**
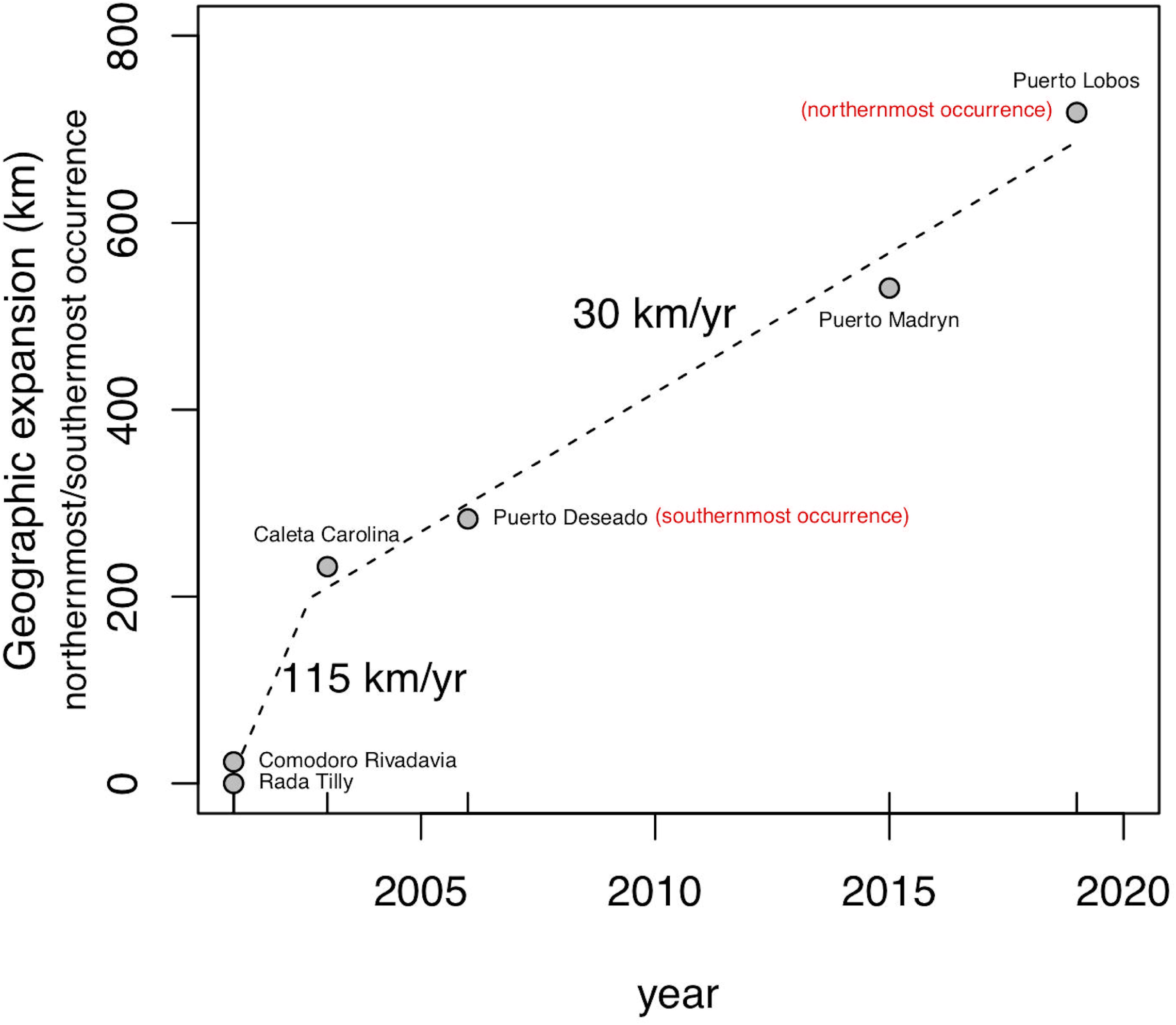
Range expansion of *C. maenas* from the likely entry point at Comodoro Rivadavia (45.46°S). The dotted line is based on a segmented regression. The expansion rates are estimated as the slope for each segment (km/yr.).

The new occurrence of *C. maenas* in Puerto Lobos fits the predictions of the ensemble SDM, which forecast a probability of occupancy higher than 60% for the area (Fig. 1b) and accurately predicted the southern sector of the San Jorge Gulf, with a probability of occupancy higher than 75% where maximum abundances have been registered at Mazaredo (Barón 2007). Our ensemble SDM (averaged across six algorithms, mean AUC = 0.96–0.99) was carried out at a much finer spatial resolution than previous studies (Compton et al. 2010), thus predicting a wider potential invasion area across the SWA. Areas with a probability of occupancy by *C. maenas* higher than 0.75 extended from southern Santa Cruz province (50°S) to southern Brazil (30°S), thereby confirming previous analyses (Compton et al. 2010). Interestingly, the southern boundary (at 47°S) has remained stable for more than 10 years, suggesting some sort of dispersal restriction (i.e., larval development at lower temperatures, de Rivera et al. 2007) and/or lack of sampling in the area.

Potential distribution of *C. maenas* predicts that this species can reach up to southern Brazil (Compton et al. 2010; this study), going through the northern San Matías Gulf, Bahía Anegada and the Bahía Blanca estuary where beds of native (*Ostrea puelchana*) an invasive (*Crassostrea gigas*) oysters are established. Recently, the presence of Ostreid herpesvirus-1 (OsHV1) in wild *C. gigas* was detected in Argentina (Barbieri et al. 2019); this virus has resulted in mass mortalities among early life stages of *C. gigas* worldwide (Burge et al. 2007; Prado-Alvárez et al. 2016). Evidence of the role of *C. maenas* in transmission dynamics of OsHV1 is now available at two Irish oyster culture sites (Bookelaar et al. 2018), while decline in oyster beds have been related to *C. maenas* predation in Atlantic Canada shorelines (Poirier et al. 2017). Hence, if *C. maenas* continues spreading northwards it would open a complete new scenario of potential predation and infection cycle through marine food webs raising concern about the conservation status of economically important oyster beds in the region.

On the other hand, it has been proposed that C*. maenas* populations from Argentina are a secondary invasion of possible Australian origin (Darling et al. 2008), so the SDM was also run using only Australian observations (n=63), and the model showed a low probability of occupancy across its current invaded range in the SWA (data not shown). However, in their genetic study, Darling et al. (2008) stated that given the limited sampling of *C. maenas* in Argentina (only one site with 15 specimens), and pending more thorough investigation of vector strength, this hypothesis of Australian origin necessitates further investigation.

Although *C. maenas* was detected nearly 20 years ago, very little information is known regarding its potential ecological impacts on coastal marine ecosystems along the SWA. *C. maenas* preyed upon native species in feeding trails (Hidalgo et al. 2007) and native kelp gulls also consumed green crabs contributing 7.3–23.9% to the overall diet (Yorio et al. 2020). In addition, most of the studies carried out have focused on intertidal habitats, while Patagonian subtidal zones remain largely unexplored (but see Battini and Bortolus 2020). Our results suggest that the pace of expansion of *C. maenas* has slowed down since its original invasion, which may give us valuable time to generate decision-making tools that will allow coastal managers to focus monitoring efforts at least on marine protected areas or zones with high concentrations of fishing resources, by designing mitigation and management plans accordingly.

## Acknowledgments

MMR’s research was funded by Fondo Nacional de Desarrollo Científico y Tecnológico (FONDECYT) under grant 1200843. SG’s research was funded by Consejo Nacional de Investigaciones Científicas y Técnicas (CONICET) under PIP-112-201701-00080.

## Figure legends

**Fig. S1.**
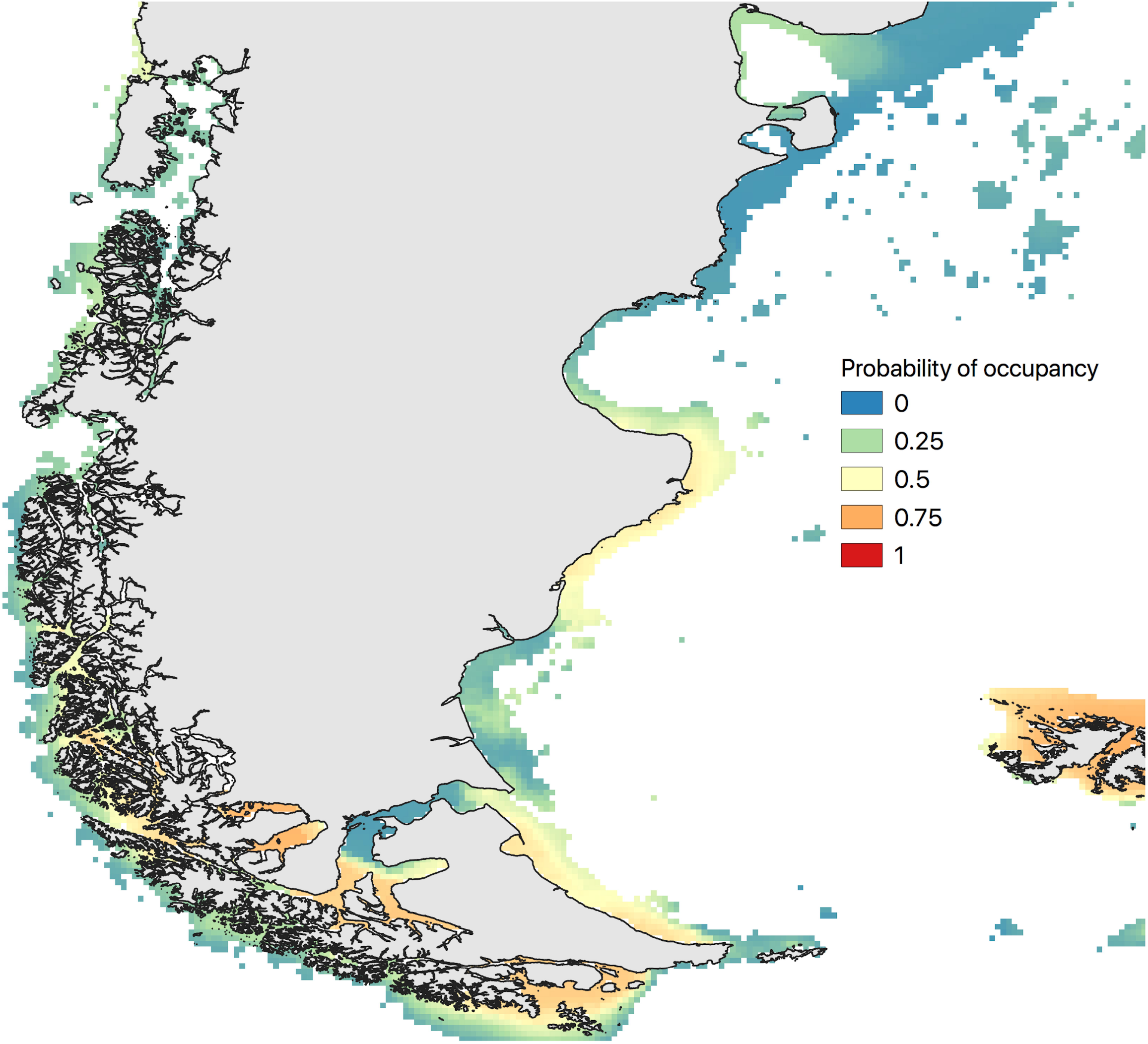
*Carcinus maenas* along the SW Atlantic coast. Species distribution model showing the invasion potential of *C. maenas*. Only Australian observations (n=63) were used.

## References

Assis J, Tyberghein L, Bosch S, Verbruggen H, Serrão EA, De Clerck O (2018) Bio-ORACLE v2. 0: Extending marine data layers for bioclimatic modelling. Glob Ecol Biogeogr. https://doi.org/10.1111/geb.12693

Barbieri ES, Medina CD, Vázquez N, Fiorito C, Martelli A, Wigdorovitz A. et al (2019) First detection of Ostreid herpesvirus 1 in wild *Crassostrea gigas* in Argentina. J. Invertebr. Pathol. https://doi.org/10.1016/j.jip.2019.107222

Barón P (2007) Introducción de especies exóticas en Patagonia: La reciente invasión del cangrejo verde europeo *Carcinus maenas* como modelo para el estudio del impacto ecológico y la planificación de estrategias de control. Proyecto PNUD ARG 02-018. Conservación de la Diversidad Biológica y Prevención de la Contaminación Marina en Patagonia. CENPAT, Puerto. Madryn.

Battini N, Bortolus A (2020) A major threat to a unique ecosystem. Front Ecol Environ. https://doi:10.1002/fee.2154

Bookelaar BE, Reilly AO, Lynch SA, Culloty SC (2018) Role of the intertidal predatory shore crab *Carcinus maenas* in transmission dynamics of ostreid herpesvirus-1 microvariant. Dis Aquat Organ. https://doi.org/10.3354/dao03264

Burge CA, Judah LR, Conquest LL et al (2007) Summer seed mortality of the Pacific oyster, *Crassostrea gigas* Thunberg grown in Tomales Bay, California, USA: the influence of oyster stock, planting time, pathogens, and environmental stressors. J Shellfish Res. https://doi.org/10.2983/0730-8000(2007)26[163:SSMOTP]2.0.CO;2

Carlton JT, Cohen AN (2003) Episodic global dispersal in shallow water marine organisms: the case history of the European shore crabs *Carcinus maenas* and *C. aestuarii*. J. Biogeogr. https://doi.org/10.1111/j.1365-2699.2003.00962.x

Compton TJ, Leathwick JR, Inglis GJ (2010) Thermogeography predicts the potential global range of the invasive European green crab (*Carcinus maenas*). Divers Distrib. https://doi.org/10.1111/j.1472-4642.2010.00644.x

Darling JA, Bagley MJ, Roman JOE, Tepolt CK, Geller JB (2008) Genetic patterns across multiple introductions of the globally invasive crab genus *Carcinus*. Mol Ecol. https://doi.org/10.1111/j.1365-294X.2008.03978.x

De Rivera CE, Hitchcock NG, Teck SJ, Steves BP, Hines AH, Ruiz GM (2007) Larval development rate predicts range expansion of an introduced crab. Mar Biol. https://doi.org/10.1007/s00227-006-0451-9

Grosholz, ED (1996) Contrasting rates of spread for introduced species in terrestrial and marine systems. Ecology. https://doi.org/10.2307/2265773

Grosholz ED, Ruiz GM (1996) Predicting the impact of introduced marine species: lessons from the multiple invasions of the European green crab *Carcinus maenas*. Biol Conserv. https://doi.org/10.1016/0006-3207(94)00018-2

Hidalgo FJ, Barón PJ, Orensanz JML (2005) A prediction come true: the green crab invades the Patagonian coast. Biol Invasions. https://doi.org/10.1007/s10530-004-5452-3

Hidalgo FJ, Silliman BR, Bazterrica MC, Bertness MD (2007) Predation on the rocky shores of Patagonia, Argentina. Estuar Coast. https://doi.org/10.1007/BF02841342

Hijmans RJ (2017) Geosphere: Spherical Trigonometry. R package version 1.5-7. https://CRAN.Rproject.org/package=geosphere. Accessed 17 March 2020

Jamieson GS, Foreman MGG, Cherniawsky JY, Levings CD (2002) European green crab (*Carcinus maenas*) dispersal: The Pacific experience. In: Paul AJ, Dawe EG, Elner R, Jamieson GS, Kruse GH, Otto S,Sainte-Marie B, Shirley TC and Woodby D (eds) Crabs in cold water regions: Biology, management and economics. 1st edn. University of Alaska Sea Grant College Program AK-SG-02-01, Alaska, pp. 561–576.

Lowe S, Browne M, Boudjelas S, De Poorter M (2000) 100 of the world’s worst invasive alien species: a selection from the global invasive species database. Invasive Species Specialist Group, Auckland.

Muggeo VM (2008) Segmented: An R package to fit regression models with broken-line relationships. RNews, No. 1, R Foundation for Statistical Computing, 20–25. http://cran.rproject.org/doc/Rnews/Rnews_2008-1.pdf. Accessed 17 March 2020

Naimi B, Araújo MB (2016) sdm: a reproducible and extensible R platform for species distribution modelling. Ecography. https://doi.org/10.1111/ecog.01881

Orensanz JML, Schwindt E, Pastorino G et al (2002) No longer the pristine confines of the world ocean: a survey of exotic marine species in the southwestern Atlantic. Biol Invasions. https://doi.org/10.1023/A:1020596916153

Pante E, Simon-Bouhet B (2013) marmap: a package for importing, plotting and analyzing bathymetric and topographic data in R. PLoS One. https://doi.org/10.1371/journal.pone.0073051

Poirier LA, Symington LA, Davidson J, St-Hilaire S, Quijón PA (2017) Exploring the decline of oyster beds in Atlantic Canada shorelines: potential effects of crab predation on American oysters (*Crassostrea virginica*). Helgol Mar Res. https://doi.org/10.1186/s10152-017-0493-z

Prado-Alvarez M, Darmody G, Hutton S et al (2016) Occurrence of OsHV-1 in *Crassostrea gigas* cultured in Ireland during an exceptionally warm summer. Selection of less susceptible oysters. Front. Physiol. https://doi.org/10.3389/fphys.2016.00492

Schwindt E (2007) The invasion of the acorn barnacle *Balanus glandula* in the south-western Atlantic 40 years later. J Mar Biol Assoc UK. https://doi.org/10.1017/S0025315407056895

Thresher R, Proctor C, Ruiz G, Gurney R, MacKinnon C, Walton W. et al (2003) Invasion dynamics of the European shore crab, *Carcinus maenas*, in Australia. Mar Biol. https://doi.org/10.1007/s00227-003-1011-1

Torres PJ, González-Pisani X (2016) Primer registro del cangrejo verde, *Carcinus maenas* (Linnaeus, 1758), en Golfo Nuevo, Argentina: un nuevo límite norte de distribución en costas patagónicas. Ecol Austral. 26:134–137.

Vinuesa JH (2005) Distribución de crustáceos decápodos y estomatópodos del golfo San Jorge, Argentina. Rev Biol Mar Oceanogr. http://dx.doi.org/10.4067/S0718-19572005000100002

Wieters EA, McQuaid C, Palomo G, Pappalardo P, Navarrete SA (2012) Biogeographical boundaries, functional group structure and diversity of rocky shore communities along the Argentinean coast. PLoS One. https://doi.org/10.1371/journal.pone.0049725

Yamada SB, Hunt C, Richmond N (2000) The arrival of the European green crab, *Carcinus maenas*, in Oregon estuaries. In: Pederson J (ed) Proceedings of the 1st International Symposium on Marine Bioinvasions. 1st edn. MIT Press, Boston, pp 24–34.

Yorio P, Suárez N, Kasinsky T, Pollicelli M, Ibarra C, Gatto A (2020) The introduced green crab (*Carcinus maenas*) as a novel food resource for the opportunistic kelp gull (*Larus dominicanus*) in Argentine Patagonia. Aquat Invasions. https://doi.org/10.3391/ai.2020.15.1.10

